## Supplementary material for "Vocal convergence in a multi-level primate society: insights into the evolution of vocal learning": Electronic Supplementary Material

Table S1: Overview over male presence and the periods during which recordings were obtained. Male ID code, Gang and party membership and recording periods are given. Orange: Mare gang; Blue: Simenti gang. Lighter shades indicate assumed presence[1], darker shades confirmed presence. Periods when recordings were taken are marked with ‘x’.

| ID | Gang | Party | 2010/11 | 2014 | 2016 |
|---|---|---|---|---|---|
| PTR | Mare | 4 | x | | |
| CSS | Mare | 4 | x | | |
| DTM | Mare | 4 | x | | |
| OSM | Mare | 4 | x | | x |
| NDR | Mare | 4 | x | x | x |
| ANT | Mare | 4 | | | x |
| AND | Mare | 9 | | x | x |
| SNE | Mare | 9 | x | x | x |
| BAA | Mare | 9 | | x | x |
| DRK | Mare | 9 | | x | x |
| VNC | Mare | 9 | | x | x |
| WNT | Mare | 9 | | | x |
| MRX | Mare | 9 | | x | x |
| MST | Simenti | 5 | x | x | x |
| ADM | Simenti | 5 | | x | x |
| FFE | Simenti | 5 | | | x |
| FDL | Simenti | 5 | | x | x |
| SPP | Simenti | 5 | | | x |
| JKY | Simenti | 6 | | x | x |
| MSA | Simenti | 6 | x | | x |
| ASN | Simenti | 6 | | | x |
| RBT | Simenti | 6 | | | x |
| MLK | Simenti | 6 | | x | x |
| BEN | Simenti | 6 | | | x |
| IND | Simenti | 6 | | x | x |
| LOU | Simenti | 6 | | | x |
| WLD | Simenti | 6 | | x | x |

[1]As Guinea baboon males are mostly philopatric, we assume that males identified as adult by 2014 were most likely already present as adolescents in 2010/11. Three subjects died during the course of the study.

Table S2. Number of calls and number of males represented per context. Calls were recorded in four major contexts: (i) affiliative interactions, including approaches and contact sitting with both males and females, (ii) greeting between males, (iii) infant handling, and (iv) looking at an interaction/resting near third parties.[1]

| Context | N calls | N males |
|---|---|---|
| Affiliation | 179 | 23 |
| Greeting | 226 | 22 |
| Infant handling | 189 | 24 |
| Look/Rest | 162 | 15 |

[1]Since previous analyses revealed only minor variation between contexts, we favored having a larger data set where males are represented by calls from multiple contexts over controlling for context.

Table S3: Acoustic parameters used by the Discriminant Function Analysis to distinguish the grunts of 27 male Guinea baboons. Variables are ordered by the correlation coefficient of the structure matrix, i.e. in relation to their importance for distinguishing between males.

| Parameter | Description |
|---|---|
| F0 mean[1)] | Fundamental frequency (F0) mean across all time segments [Hz] |
| PF max[1)] | Maximum PF of all time segments (peak frequency: highest frequency amplitude of a time segment) [Hz] |
| DFA3 end[2)] | End DFA 3rd quartile (DFA: distribution of frequency amplitude) [Hz] |
| DFA3 med[2)] | Median DFA 3rd quartile [Hz] |
| F0 slope[1)] | Factor of linear trend of F0 |
| DFA3 start[2)] | Start DFA 3rd quartile [Hz] |
| DFA3 maloc[2)] | Location of maximum DFA 3rd quartile [(1/duration)*location]) |
| DFA3 min[2)] | Minimum DFA 3rd quartile [Hz] |
| Corr mean[2)] | Mean correlation coefficient of successive time segments |
| F1 mean[2)] | Mean frequency of 1st general amplitude peak [Hz] |
| Corr maloc[2)] | Location of maximum correlation coefficient [(1/duration)*location]) |
| Duration[2)] | Duration of call [ms] |
| PF minloc[2)] | Location of minimum PF [(1/duration)*location]) |
| DFA2 med[2)] | Median DFA 2nd quartile [Hz] |
| F1w max[2)] | Maximum frequency range of 1st general amplitude peak [Hz] |
| Duration[1)] | Duration of call [ms] |
| Noise mean[1)] | Mean noisiness measured as Wiener entropy [0-1] |
| Noise max[1)] | Maximum noisiness measured as Wiener entropy [0-1] |
| PF min[2)] | PF minimum [Hz] |
| PF maloc[2)] | Location of maximum PF [(1/duration)*location]) |
| FP1a max[2)] | Maximum amplitude of 1st general amplitude peak [rel. amplitude] |
| DFA3 max[2)] | Maximum DFA 3rd quartile [Hz] |
| DFA3 mean[2)] | Mean DFA 3rd quartile [Hz] |
| DFA2 start[2)] | Start DFA 2nd quartile [Hz] |
| FP1a mean[2)] | Mean amplitude of 1st general amplitude peak [rel. amplitude] |
| F2 mean[2)] | Mean frequency of 2nd general amplitude peak [Hz] |
| F1w end[2)] | End frequency range of 1st general amplitude peak [Hz] |
| F1w min[2)] | Minimum frequency range of 1st general amplitude peak [Hz] |
| Freq range min[2)] | Mean frequency range across all time segments [Hz] |
| F2 %[2)] | Percentage of 2nd general amplitude peak [Hz] |
| F1w mean[2)] | Mean frequency range of 1st general amplitude peak [Hz] |

[1)] Estimated from frequency time spectra with a frequency range of 500 Hz; [2)] Estimated from frequency time spectra with a frequency range of 2500 Hz.

Table S4. Correlation coefficients (Spearman) between simulated relatives and genetic relatedness estimates. The Wang estimator showed the second highest correlation coefficient, only slightly worse than Trio-ML. Due to its distribution from -1 to 1, it is better suited for correlation analyses (with acoustic similarity), and was therefore selected as the relatedness estimator. See refs. [27-29] for details.

| Estimator | Correlation Coefficient |
|---|---|
| TRIOML: | 0.847 |
| WANG: | 0.844 |
| LYNCHLI: | 0.837 |
| LYNCHRD: | 0.812 |
| RITLAND: | 0.771 |
| QUELLERGT: | 0.839 |
| DYADML: | 0.843 |

Table S5. Output of permuted DFA based on the procedure described in Mundry & Sommer 2007 (Ref. 24).

| | |
|---|---|
| "no.corr.classified.selected" | 107.37 |
| "expected.no.corr.classified.selected" | 93.83337 |
| "percent.correctly.classified.selected" | 79.5333333333333 |
| "expected.percent.correctly.classified.selected" | 69.5062 |
| "P.for.selected" | 0.006 |
| "no.corr.cross.classified" | 69.67 |
| "expected.no.corr.cross.classified" | 23.11867 |
| "percent.corr.cross.classified" | 11.219001610306 |
| "expected.percent.corr.cross.classified" | 3.72281320450886 |
| "P.for.cross.classified" | 0.001 |
| "no.levels.contr.factor" | 756 |
| "no.levels.contr.factor.selected" | 135 |
| "no.levels.test.factor" | 27 |
| "no.cases" | 756 |
| "no.cases.selected" | 135 |
| "no.cases.selected.per.level.of.control.fac" | 1 |
| "no.permutations" | 1000 |
| "no.random.selections" | 100 |

Table S6: Acoustic parameters used to distinguish between gangs and parties. Variables are ordered by the correlation coefficient of the structure matrix, i.e. in relation to their importance for distinguishing between social levels. For the explanation of the variables, see Table S2.

Between gangs

| Parameter | correlation |
|---|---|
| DFA3 end | -0.51 |
| DFA2 maloc | -0.45 |
| DFA3 maloc | -0.42 |
| DFA2 end | -0.4 |
| DFA3 max | -0.31 |
| DFA2 max | -0.26 |

Between parties

| Parameter | correlation |
|---|---|
| DFA3 end | -0.49 |
| DFA2 maloc | -0.43 |
| DFA3 maloc | -0.38 |
| DFA2 end | -0.37 |
| DFA3 max | -0.27 |
| F0 maloc | -0.26 |
